# Do calcium channel blockers applied to cardiomyocytes cause increased channel expression resulting in reduced efficacy?

**DOI:** 10.1101/2023.10.09.561500

**Authors:** Karoline Horgmo Jæger, Verena Charwat, Samuel Wall, Kevin E. Healy, Aslak Tveito

## Abstract

In the initial hours following the application of the calcium channel blocker (CCB) nifedipine to microtissues consisting of human induced pluripotent stem cell-derived cardiomyocytes (hiPSC-CMs), we observe notable variations in the drug’s efficacy. Here, we investigate the possibility that these temporal changes in CCB effects are associated with adaptations in the expression of calcium ion channels in cardiomyocyte membranes. To explore this, we employ a recently developed mathematical model that delineates the regulation of calcium ion channel expression by intracellular calcium concentrations. According to the model, a decline in intracellular calcium levels below a certain target level triggers an upregulation of calcium ion channels. Such an upregulation, if instigated by a CCB, would then counteract the drug’s inhibitory effect on calcium currents. We assess this hypothesis using time-dependent measurements of hiPSC-CMs dynamics and by refining an existing mathematical model of myocyte action potentials incorporating the dynamic nature of the number of calcium ion channels. The revised model forecasts that the CCB-induced reduction in intracellular calcium concentrations leads to a subsequent increase in calcium ion channel expression, thereby attenuating the drug’s overall efficacy. The data and fit models suggests that dynamic changes in cardiac cells in the presence of CCBs may be explainable by induced changes in protein expression, and that this may lead to challenges in understanding calcium based drug effects on the heart unless timings of applications are carefully considered.

## 1 Introduction

Excitable cells exhibit electrochemical homeostasis over extended periods, even though the constituent membrane proteins (e.g., ion channels), which underpin the action potential of these cells, are in a continuous state of renewal. How is the expression of ion channels controlled to preserve the electrical properties essential for physiological functions? This pressing question has been the subject of extensive investigation by Marder and colleagues, among others, over several years; see, e.g., [1, 2, 3, 4, 5, 6, 7].

In one significant contribution, O’Leary et al. [4] developed a mathematical framework to represent ion channel expression. Their work was founded on the hypothesis that the intracellular calcium concentration governs ion channel expression levels. According to their model, if the calcium concentration falls below a specific target value, the number of ion channels carrying calcium ions will, through a multistep process of DNA transcription, RNA translation, and protein trafficking, increase until that target value is met.

Their modeling framework has found applications beyond the original scope. Recently, Moise and Weinberg [8] adapted this model to analyze ion conductances within the cells of the Sinoatrial node, further demonstrating the model’s relevance and utility in understanding the complex dynamics of ion channel regulation; see also [9].

### Modeling expression of calcium channels

One version of the model for dynamic ion channel expression can be written in the following form,

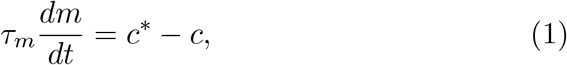

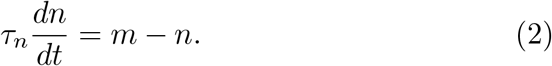

A description of how the model from [4, 8] can be rewritten to the form (1)–(2) is provided in the Supplementary Information (SI). Here, *τ*_*m*_ and *τ*_*n*_ are time constants, *c* is the cytosolic calcium concentration, and *c*\* is the target level of the cytosolic calcium concentration. Furthermore, we let *M*_0_ and *N*_0_ represent the default numbers of messenger RNAs (mRNAs) and the number of expressed calcium ion channel proteins, respectively. The associated dynamic numbers are given by *M* (*t*) = *m*(*t*)*M*_0_ and *N* (*t*) = *n*(*t*)*N*_0_, where the evolution of the relative changes *m* and *n* are governed by the system (1)–(2). The associated calcium current is given by

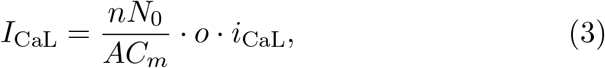

where *A* is the area of the cell membrane (in *μ*m^2^), *C*_*m*_ is the specific membrane capacitance (in pF/*μ*m^2^), *o* is the open probability, and *i*_CaL_ is the average current through a single open calcium channel (in pA). When a CCB is present, we assume that the current through a single calcium channel is reduced. We incorporate this in the model by introducing a scaling factor *b*(*D*) corresponding to the reduction in single-channel current

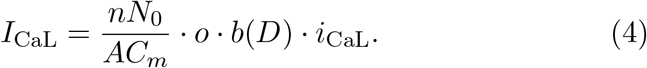

When no drug is present *b*(0) = 1, and when a drug blocks the current by, for instance, 90%, then *b*(*D*) = 0.1.

In Figure 1, the properties of this model are demonstrated in conjunction with a mathematical model of the action potential for hiPSC-CMs (see [10] and the SI for details). Initially, we set *n* = *m* = 1*/*10 and the initial conditions for the remaining model variables (including *c*) are found by running a simulation of the model with *n* fixed at 0.1. The parameter *c*\* is set as the average cytosolic calcium concentration during an action potential (AP) cycle in the default model with *n* = 1. Furthermore, we set *τ*_*m*_ = 400 mMms and *τ*_*n*_ = 1000 ms. The model forecasts a significant upregulation in the number of calcium channels, leading to a normalization of the intracellular calcium concentration. After a span of 20 hours, the ion channel count is fully restored, and the intracellular calcium concentration *c* attains its target level *c*\* on average during an AP cycle. Notably, this membrane rectification process unfolds over hours, contrasting with the brief duration of each action potential, which lasts between 200–600 ms. This example underscores the model’s utility: if the intracellular calcium concentration diverges from the target level, the ion channel count will be modulated to bring the concentration back to equilibrium. This aligns with the model’s original intent, as corroborated by studies such as [3, 4].

**Figure 1.**
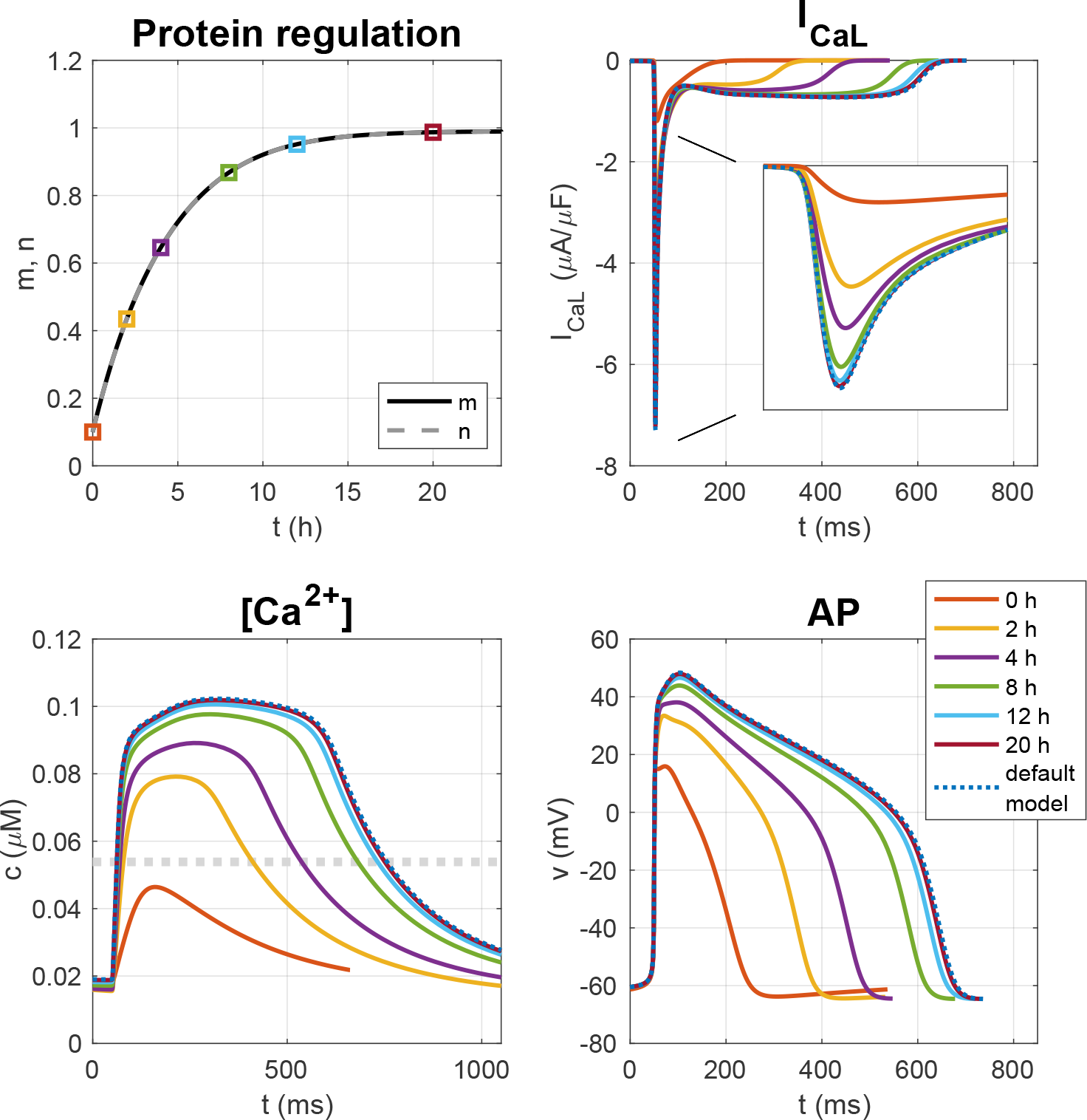
Simulation of the system (1)–(2) included in an AP model for hiPSC-CMs with *τ*_*m*_ = 400 mMms and *τ*_*n*_ = 1000 ms from a starting point of *n* = *m* = 0.1. The upper left plot shows the time evolution of *m* and *n*. In addition, six time points are marked. The calcium current density over the cell membrane, *I*_CaL_, the cytosolic calcium concentration, *c*, and the action potential (AP) at these six points in time are plotted in the next three figure panels. In addition, we show the default model solution, and we observe that after 20 h of simulation, *m* and *n* are very close to 1 and the model solution is very close to the default model solution. The dotted gray line in the lower left plot shows the target calcium concentration, *c*\*.

### Consequences for calcium channel blockers

CCBs are a class of drugs that have wide clinical cardiovascular applicability, including the treatment of hypertension, angina, and cardiac arrhythmias [11]. Through L-type calcium antagonism, they can have an effect on both vascular smooth muscle and myocardial muscle, and can cause the reduction of blood pressure, coronary-artery dilation, and depression of cardiac contractility [12]. In the heart, the block of *I*_CaL_ diminishes the influx of calcium into the intracellular space during an AP. This, in turn, results in a decreased release of calcium from internal storage units, because of the effect commonly known as graded release (see, e.g., [13, 14, 15]). Consequently, the average intracellular calcium concentration should be reduced when exposed to CCBs. In the presented framework, when the intracellular calcium concentration *c* falls below the target value *c*\*, the model predicts an upregulation in the expression of calcium channels, thereby increasing the calcium influx into the cell. If this trend continues unchecked, the inhibitory effect of the CCB may therefore be nullified.

An illustration of the model is provided in the left panel of Figure 2. In this simulation, the calcium current is reduced by 90% (i.e., *b*(*D*) = 0.1 in (3)). As the blocking lowers the average cytosolic calcium concentration, the model anticipates an increase in the number of ion channels, eventually counteracting the blocking effect; the calcium target is met, and the current strength is completely restored.

**Figure 2.**
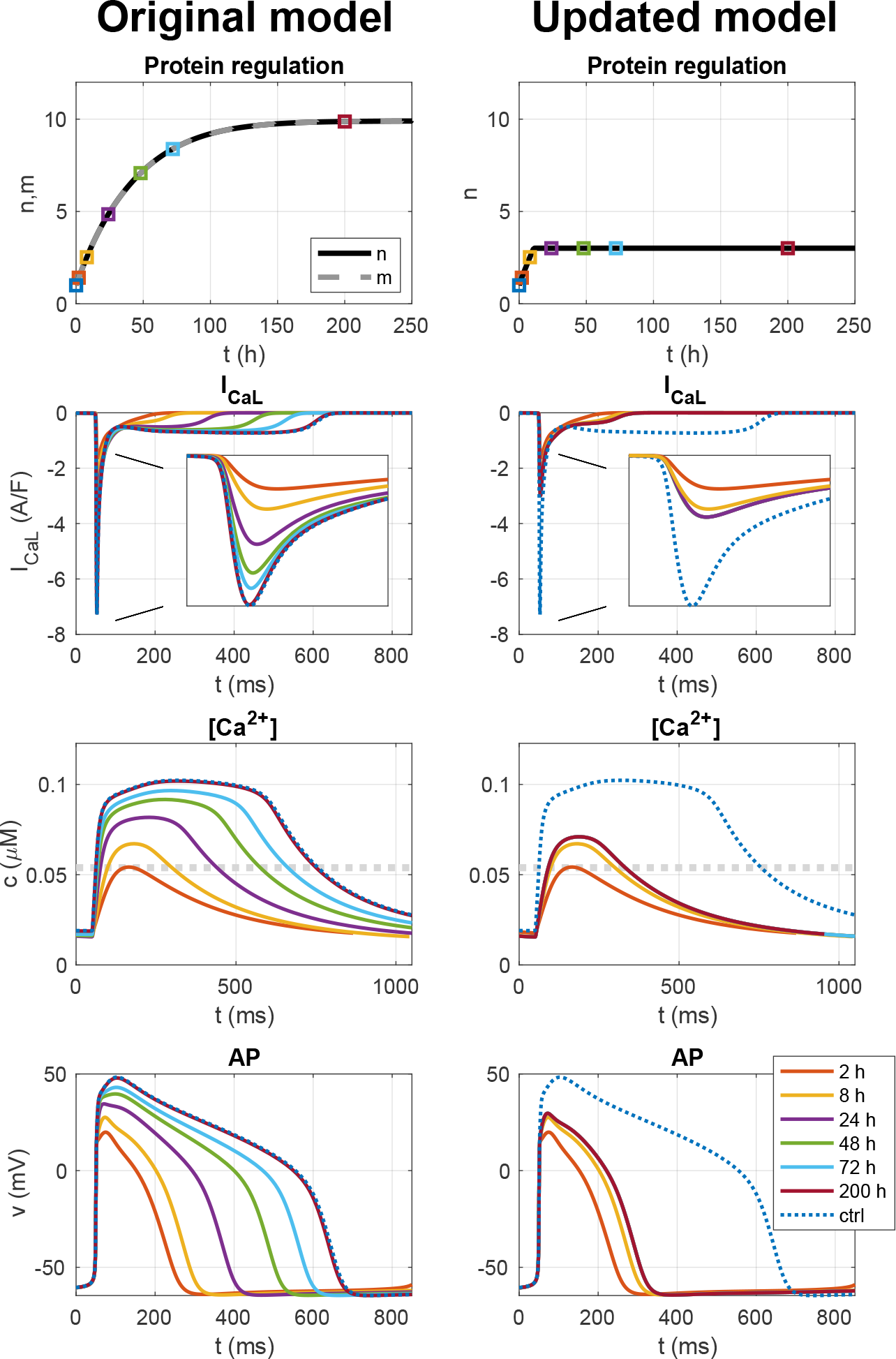
Left panel: Simulation of the system (1)–(2) included in an AP model for hiPSC-CMs with *τ*_*m*_ = 400 mMms and *τ*_*n*_ = 1000 ms from a starting point of *n* = *m* = 1 and a 90% block of *I*_CaL_ (*b*(*D*) = 0.1). Initially, *c* is considerably reduced by the *I*_CaL_ block, leading to an increase in *m* and *n*. Eventually, *m* and *n* reach a value of 10, restoring *I*_CaL_ to its default strength as well as restoring the cytosolic calcium concentration and the AP to the default (control, ctrl) case. Right panel: Simulation of the same case using the updated model (13) with *τ*_*n*_ = 400 mMms, *n*_*−*_ = 0.1 and *n*_+_ = 3. In this case, *n* cannot increase above *n*_+_, and the model reaches an equilibrium solution at *n* ≈ *n*_+_ and the blocking of *I*_CaL_ is not completely diminished over time.

This example highlights the need for model refinement when considering the presence of CCBs; unlimited growth in the number of ion channels is not realistic. In the following sections, we demonstrate that the model can be simplified from a 2 × 2 system to a scalar equation, imposing a limit on ion channel growth. Additionally, we show that the model’s predictions align well with experimental observations using hiPSC-CMs. In summary, both the mathematical model and the *in vitro* measurements indicate a time-dependent effect of the CCB. The blocking effect is most pronounced in the initial hours but gradually diminishes, without being completely nullified, consistent with the theory that channel expression will increase when calcium concentrations fall below the target level.

## 2 Methods

We will address the effect of CCBs using a mathematical model motivated by the system (1)–(2). However, since *m* and *n* are very similar, we will show that the 2 × 2 system can be faithfully replaced by a scalar equation. Furthermore, we will update the model in order avoid unlimited growth or reduction in the gene expression. We will also describe how the effect of the CCB is measured using hiPSC-CMs.

### 2.1 Reduction to a scalar model

Both in Figure 1 and in Figure 2 we noticed that *m* ≈ *n*. This motivates analysis of the deviation given by

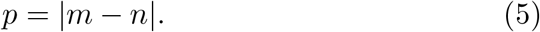

From the system (1)–(2) we get,

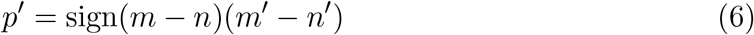

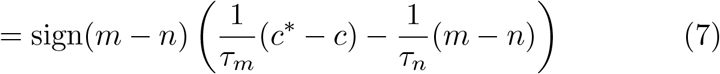

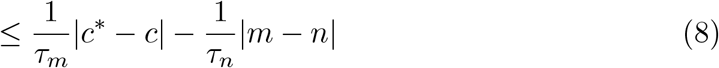

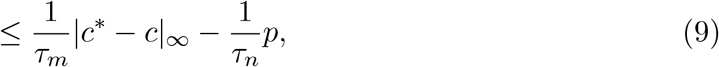

where |*c*\* *−c*|_*∞*_ denotes the largest deviation of *c* from the target value *c*\*. From Gronwall’s inequality ([16], page 283) we get

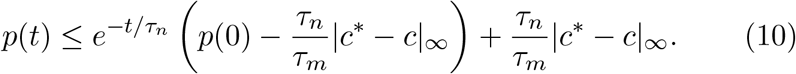

Since *τ*_*n*_ is on the order of a few seconds or less, we see that for large values of *t* (hours), we have

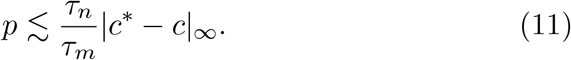

Numerical experiments reveals that in order to fit data from measurements of hiPSC-CMs (see below), we need the parameter *τ*_*m*_ to be around 400 mM×ms. Furthermore, varying *τ*_*n*_ between 10 ms and 10, 000 ms did not seem to influence the main results (see the SI). Moreover, we observe that |*c*\* *− c*|_*∞*_ never exceeds 5 × 10^*−*5^mM. Therefore, we have

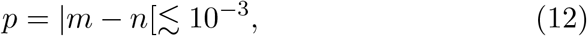

so *m* ≈ *n* and we can reduce the system to a scalar equation in *n*.

### 2.2 Introducing upper and lower bounds on the protein expression

We have seen that the original model runs into difficulties when the calcium current is blocked. The source of the difficulty is that the number of ion channels are allowed to grow in an unlimited manner. This is not realistic, and we want to adjust the model accordingly by putting an upper limit, *n* = *n*_+_, on how much the number calcium ion channels can grow. In addition, we wish to enforce a lower limit, *n* = *n*_*−*_ so that the number of ion channels cannot become negative. To this end, we introduce the model

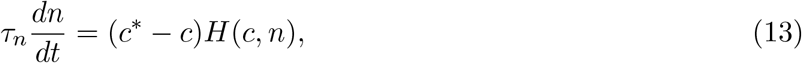

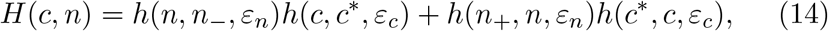

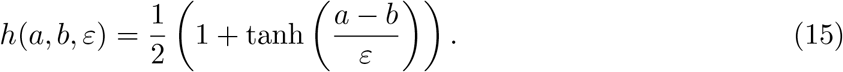

Here, the parameters *τ*_*n*_, *c*\* and *n*_+_ must be estimated, and we let *n*_*−*_ = 0.1, *ε*_*n*_ = 0.01 and *ε*_*c*_ = 10^*−*7^ mM. Note that *n* is a dimensionless number and *nN*_0_ is the total number of ion channels in the membrane. The unit of concentrations are mM and time is in ms, hence the unit of *τ*_*n*_ is ms×mM. The function *H* is introduced to bound the number of ion channels and is illustrated in Figure 3.

**Figure 3.**
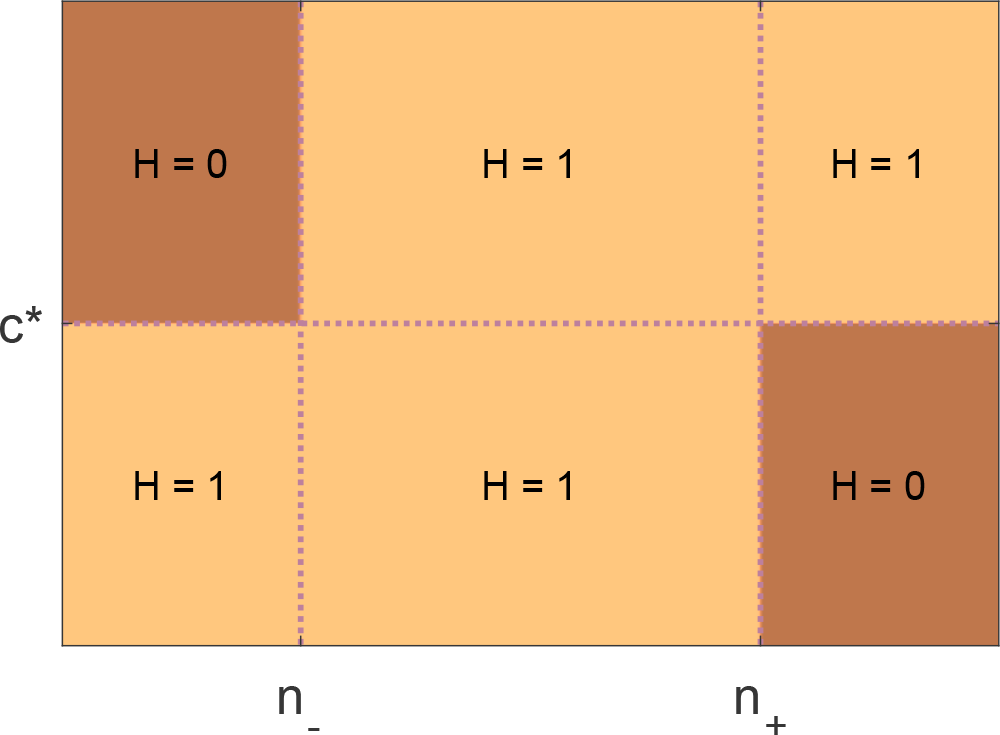
Illustration of the function *H* defined in (14)–(15). The function is 1 everywhere except that it reduces to 0 when *n < n*_*−*_ and *c > c*\* and when *n > n*^+^ and *c < c*\*. The steepness of the transition between the different values of *H* is controlled by the *ε*_*n*_ and *ε*_*c*_ parameters.

The model (1)–(2) reaches its equilibrium when the calcium concentration reaches the target value, i.e., when *c* ≈ *c*\*. The new model (13) can reach an equilibrium state either if the target value is achieved, if the maximum number of calcium ion channels is reached, i.e. if *n* ≈ *n*_+_, or if the minimum number of calcium ion channels is reached, i.e. if *n* ≈ *n*_*−*_.

In the right panel of Figure 2, we have rerun the *I*_CaL_ block example illustrated in the left panel of Figure 2 using the updated version of the model (i.e., (13)). We use the parameters *τ*_*n*_ = 400 mMms and *n*_+_ = 3. Moreover, *c*\* is still the average cytosolic calcium concentration during an AP cycle in the default model with *n* = 1. In the right panel of Figure 2, we observe that in the new model, the number of calcium channels, *n*, initially increases in response to the reduced calcium concentration resulting from the reduction of *I*_CaL_. However, when *n* reaches the value of *n* = *n*_+_ = 3, *n* is not able to increase further and reaches an equilibrium solution even though the average cytosolic calcium concentration has not reached *c*\*. In other words, the effect of the *I*_CaL_ block is reduced over the initial hours after *I*_CaL_ block, but *I*_CaL_ is not able to fully recover as in the left panel for the original model (1)–(2).

### 2.3 Measuring the effect of CCB in hiPSC-CMs

Cardiac microtissues were generated based on a refined version of our previously published cardiac microtissue platform [17, 18]. In brief, microtissues were created in 384 well plates (206384; GraceBioLabs) with custom substrates (COC TPE E140; Stratec) featuring 4 tissue formation chambers per well (1400x200x150 *μ*m length×width×depth) using 35000 iPSC-derived cardiomyocytes (C1056; Fujifilm CDI) per well. Microtissues were used at day 29 and stained with 500nM voltage sensitive dye (PhoS1; Photoswitch Biosciences). Nifedipine (PHR1290; Sigma) was freshly prepared as 6-fold concentrate in cell culture media (M1003; Fujifilm CDI) from a 10mM stock in DMSO. After baseline recordings, 10 *μ*L of concentrated drug were added to 50 *μ*L culture media in each well to achieve the desired dose of 0.1 or 1 *μ*M. Repeated recordings were performed at 2, 4, 6, 8, 13 and 16h following drug addition. All imaging was performed inside the incubation chamber of an ImageXpress Micro (Molecular Devices) microscope. Cy-5 fluorescence (1.5% laser power) was recorded in each well for 8s at 50fps via a 4x objective with 2x2 binning. Each video was then segmented to obtain voltage traces for individual tissues and biomarkers (APD50, APD80 and beat rate) were computed.

## 3 Results

In this section, we will illustrate that the model (13) coupled to a model for the action potential of hiPSC-CMs is able to qualitatively represent the temporal changes observed in the effect of the CCB nifedipine on a collection of hiPSC-CMs.

The parameters of the model are set to *τ*_*n*_ = 400 mMms and *n* = *n*_+_ = 3, as these values appeared to make the model behave relatively similar to the measurements of hiPSC-CMs. We consider two doses of nifedipine, 0.1 *μ*M and 1 *μ*M. The effect of 0.1 *μ*M nifedipine was modeled by reducing *I*_CaL_ by 55% whereas the effect of 1 *μ*M of nifedipine was modeled by reducing *I*_CaL_ by 88%. In other words, *b*(0.1 *μ*M) = 0.45 and *b*(1 *μ*M) = 0.12 in (3). These blocking percentages are in relatively good agreement with measurements of the drug effect from literature (see the SI).

### 3.1 Simulation and measurements of the effect of 0.1 *μ*M of nifedipine on hiPSC-CMs

Figure 4 shows the results of the simulations of the two doses of nifedipine as well as biomarker values collected from measurements of hiPSC-CMs. In Figure 4A, 0.1 *μ*M of nifedipine is applied. In the simulation results reported in the upper panel, we observe that the reduction of *I*_CaL_ cause a reduced cytosolic calcium concentration, which leads to a gradual increase in the number of calcium channels, *n*. Furthermore, after 16 h of drug exposure, the number of calcium channels has increased so much that *I*_CaL_, the calcium concentration and the AP is almost completely recovered to the control case. In the second row of Figure 4A, three biomarkers are computed in the control case before drug exposure and after 2 h, 4 h, 6 h, 8 h, 13 h and 16 h of drug exposure. The open gray circles are computed from the model solution and the filled colored circles are computed from optical measurements of hiPSC-CMs. For both the model and in the measurements of hiPSC-CMs, we clearly observe that the initial decrease in APD and increase in beat rate gradually is reduced over the 16 h of drug exposure. Furthermore, both the measured data and the model results suggest that the drug effect has almost completely diminished after 16 h.

**Figure 4.**
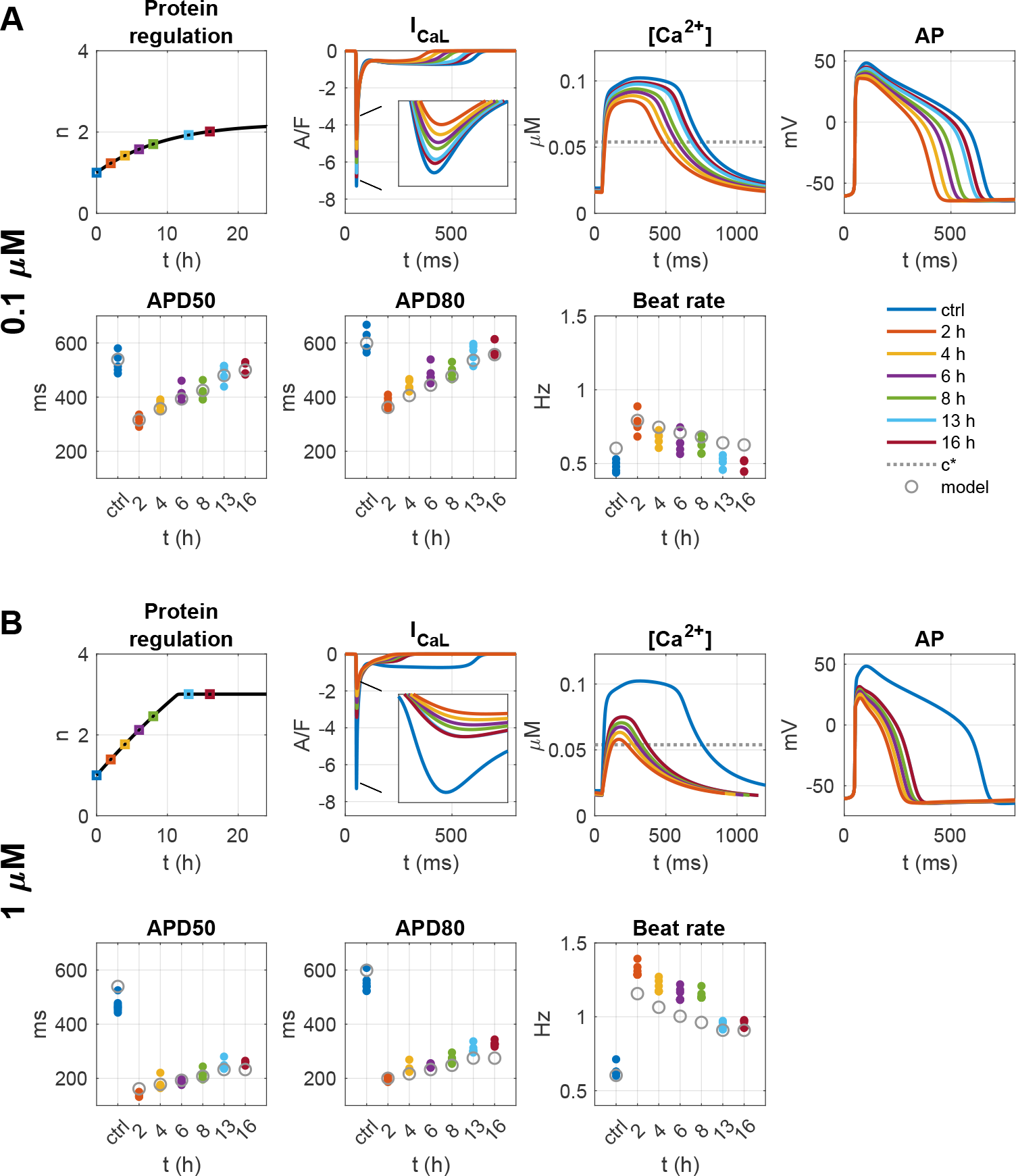
Simulated and measured effect of two doses (0.1 *μ*M and 1 *μ*M) of nifedipine on spontaneously beating hiPSC-CMs. The first and third rows of plots show properties of the model solution for the 0.1 *μ*M and 1 *μ*M doses, respectively. More specifically, the leftmost plots show the time evolution of the number of calcium channels, *n*. The next plots show *I*_CaL_, the cytosolic calcium concentration and the action potential measured in the control case (ctrl, no *I*_CaL_ block) and at six time points after the *I*_CaL_ block was applied. Furthermore, the second and fourth rows show the APD50, APD80 and beat rate sampled in the control case and at the six points in time. The open gray circles are computed from the model solution and the filled colored circles are measurements of hiPSC-CMs in the control case (blue) and after exposure to nifedipine. In the model, 0.1 *μ*M and 1 *μ*M of nifedipine were represented by reducing *I*_CaL_ by 55% and 88%, respectively, i.e. *b*(0.1 *μ*M) = 0.45 and *b*(1 *μ*M) = 0.12 in (3).

### 3.2 Simulation and measurements of the effect of 1 *μ*M of nifedipine on hiPSC-CMs

In Figure 4B, 1 *μ*M of nifedipine is applied. Again, we observe an initial decrease in cytosolic calcium concentration resulting in a gradual increase in the number of calcium channels, *n*. However, in this case, the maximum number of calcium channels, *n*_+_ is in this case reached before *I*_CaL_ is fully recovered and the increase in *n* halts. Consequently, even after 16 h of drug exposure the effect of the drug is still considerable, although somewhat less pronounced than after only 2 h of drug exposure.

## 4 Discussion

A series of recent papers propose that the gene expression of membrane proteins in excitable cells is regulated by the intracellular calcium concentration *c*, relative to a target concentration *c*\*, see, e.g., [3, 4, 8]. Simply put, if *c < c*\*, the cell compensates by increasing the number of ion channels responsible for the calcium current, aiming to elevate *c* to *c*\*. This hypothesis has been formulated through ordinary differential equations, which can be integrated into existing mathematical models of cellular action potentials.

Here, we consider the implications of this theory when a calcium channel blocker (CCB) is introduced. The application of a CCB leads to a reduction in calcium influx, resulting in *c < c*\*. According to the theory, this will trigger an increase in the number of calcium ion channels. While each individual channel’s blocking by the CCB remains constant, the overall cellular calcium current would increase due to the greater number of channels, changing the CCB’s effect. The theory suggests this process would continue until *c* = *c*\*, a notion we find implausible given the well-documented efficacy of CCBs in reducing calcium currents. To address this, we have refined the mathematical model by imposing limits on the minimum and maximum number of ion channels.

Our updated model aligns reasonably well with empirical data as illustrated in Figure 4. The model predicts that the CCB’s impact is most pronounced immediately after administration, gradually diminishing over subsequent hours. Importantly, the model changes implemented allows the CCB’s effect to not vanish but to stabilize at a significant level consistent with experimental observations.

### Timing of measurements of efficacy

Our findings suggest that the efficacy of CCBs on whole-cell currents on cardiac cells should be evaluated over an extended timeframe, rather than solely immediately post-administration. This is as dynamic changes to the calcium control system may confound correct interpretation of drug effects. We do not anticipate any temporal variations in the efficacy of CCBs at the single-channel level; the observed temporal effects are only attributed to changes in gene expression induced by changes in intracellular calcium concentrations.

### Effect on other ion channels and calcium regulatory processes

Previous studies (e.g., [4] and [8]) propose that intracellular calcium concentrations also modulate other membrane currents, regulatory systems and even intracellular calcium storage systems. Our focus here has been solely on the L-type calcium current to explore this mechanism as a description of experimental data. While other channels and regulatory processes may be changing concurrently in the system to adjust the overall calcium and membrane dynamics, we currently lack the data to evaluate broader *c* and *c*\* calcium homeostasis mechanisms.

### Clinical implications

Calcium Channel Blockers (CCBs) are widely used in clinical practice, as evidenced by multiple studies, see e.g., [19, 20]. Given the well-characterized, long-term applications of CCBs in clinical settings, our findings are unlikely to cause changes in clinical practice. However, it’s worth noting that the initial potency of CCBs may significantly exceed their long-term effects, a nuance that could be relevant for future research and treatment strategies. It is also worth noting that if calcium homeostasis mechanisms can alter the transcriptome of key ion channels, this phenomena may have an important consideration in processes that depend on the delicate balance of membrane ion channels, such as the generation of cardiac arrhythmias.

### Changes in transcriptome caused by CCBs

While our study employs mathematical models to analyze gene expression changes and utilizes calcium-sensitive dyes to measure the effects of Calcium Channel Blockers (CCBs) on hiPSC-CMs, other researchers have directly measured the cardiomyocyte transcriptome following CCB treatment. In [21], it was observed that *“CCB can lose its inhibitory effect on L-type calcium channels after chronic treatment in some iPSC-CM lines*.*”* Furthermore, the authors add that *“While acute treatment of CCBs exerted expected negative chronotropic and inotropic effects in all lines, we observed line-specific recovery after long-term treatment in two lines. Electrophysiological study showed that the calcium current was no longer inhibited in those lines after chronic treatment*.*”* In sum, the combined results motivates further analysis of time-dependent effect of CCB and other drugs affecting the membrane of cardiomyocytes.

## Acknowledgements

This work was supported by the Research Council of Norway via INPART grant agreement #322312 (SIMBER).

## Author contributions

K.H.J and A.T developed the computational methodology, designed the numerical experiments and wrote the main manuscript text. K.H.J. wrote the simulation code. A.T. conceived the project. S.W. and V.C. performed measurements of drug effects on hiPSC-CMs. K.E.H supervised experimental procedures. All authors reviewed and approved final manuscript.

## Additional information

### Disclosure of writing assistance

During the preparation of this manuscript, the authors utilized the ChatGPT4 language model to enhance the language quality for contributions from non-native English speakers. Subsequent to this automated assistance, the authors rigorously reviewed and edited the manuscript to ensure its accuracy and integrity. The authors assume full responsibility for the content of the publication.

## Data availability

The data and code generated in this study are publicly available at Zenodo: [link to be included].

## Supplementary Information

### S1 Reformulating the protein regulation model from [4, 8]

In [4, 8], the protein regulation model is given in the form^1^

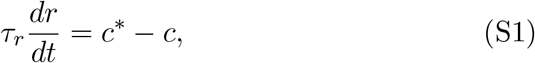

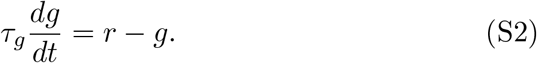

Here, *c* is the cytosolic calcium concentration, *c*\* is the target calcium concentration, *g* is the conductance of the considered type of ion channel, and *τ*_*r*_ and *τ*_*g*_ are time constants.

In our computations, we have rewritten the original formulation (S1)–(S2) to instead of *r* and *g*, model unitless scaling factors *n* and *m* for the number of ion channels in the cell membrane and the number of messenger RNAs (mRNAs). More specifically, the number of ion channels and the number of mRNAs are given by *N* (*t*) = *n*(*t*)*N*_0_ and *M* (*t*) = *m*(*t*)*M*_0_, respectively, where *N*_0_ and *M*_0_ are the default number of channels and mRNAs.

The total current through all the *I*_CaL_ channels in the cell membrane can be expressed as

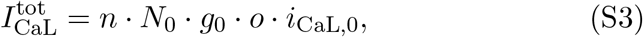

where *g*_0_ is the single channel conductance of the channel, *o* is the open probability of the calcium channels and *i*_CaL,0_ is an expression for how the single-channel current depends on model variables like the membrane potential and the cytosolic calcium concentration. In this setting, the conductance *g* in the model (S1)–(S2) is given by

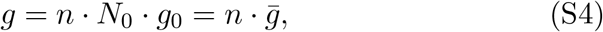

where 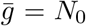 *g*_0_ is the default value of · *g* (corresponding to *n* = 1). In other words, we have

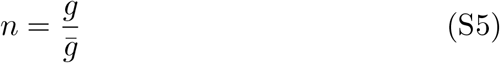

In order to rewrite (S1)–(S2) to a system for *n* and *m*, we similarly define

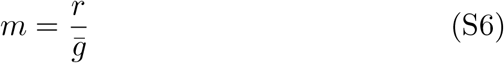

and divide both (S1) and (S2) by 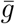. This yields

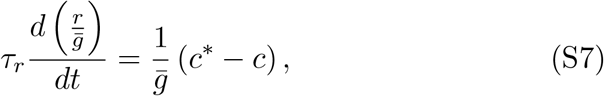

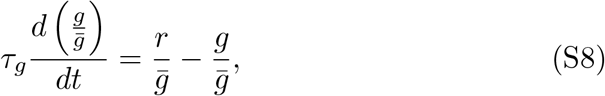

which can be rewritten as

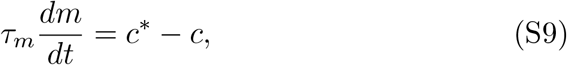

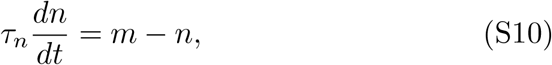

where

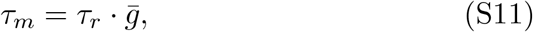

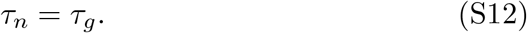

### S2 Investigating the effect of adjusting *τ*_*n*_

In Figure S1, we show the time evolution of *m* and *n* in a simulation with the model (S9)–(S10) coupled to the action potential model of hiPSC-CMs described in Section S4. The simulation is started from *n* = *m* = 0.1 in exactly the same manner as in Figure 1 in the paper. We let *τ*_*m*_ = 400 mMms (like in the paper) and vary *τ*_*n*_ between 100 ms and 10,000 ms. For comparison, the values *τ*_*n*_ = 1000 ms was used in [8]. We observe that all the considered values of *τ*_*n*_ provide almost identical solutions.

**Figure S1:**
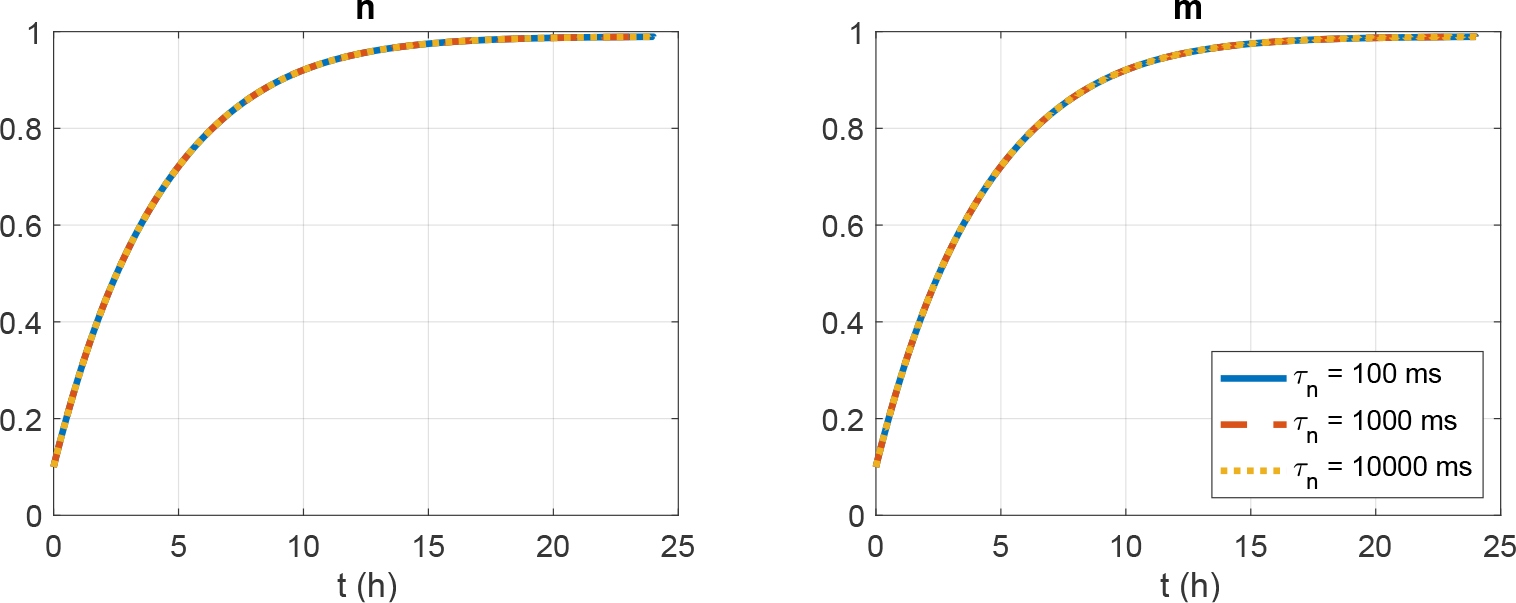
Time evolution of the solutions *n* and *m* of the model (S9)–(S10) coupled to an action potential model of hiPSC-CMs for different values of *τ*_*m*_. The simulations are started from *n* = *m* = 0.1 and *τ*_*m*_ = 400 mMms.

### S3 Comparison of nifedipine block percentages from literature

In order to compare the assumed block percentages associated with 0.1 *μ*M and 1 *μ*M of nifedipine, we identified dose-dependent effects on *I*_CaL_ from literature. The selected block percentages are compared to the data from literature in Figure S2. The figure shows the value of *b*(*D*) as a function of the dose *D* reported in different studies in addition to the values used in this study. More specifically, the effect of the drug is typically reported in terms on a half-maximal effective concentration, EC_50_, a Hill coefficient, *h* and a maximal effect *E*, and *b*(*D*) is assumed to be given by

**Figure S2:**
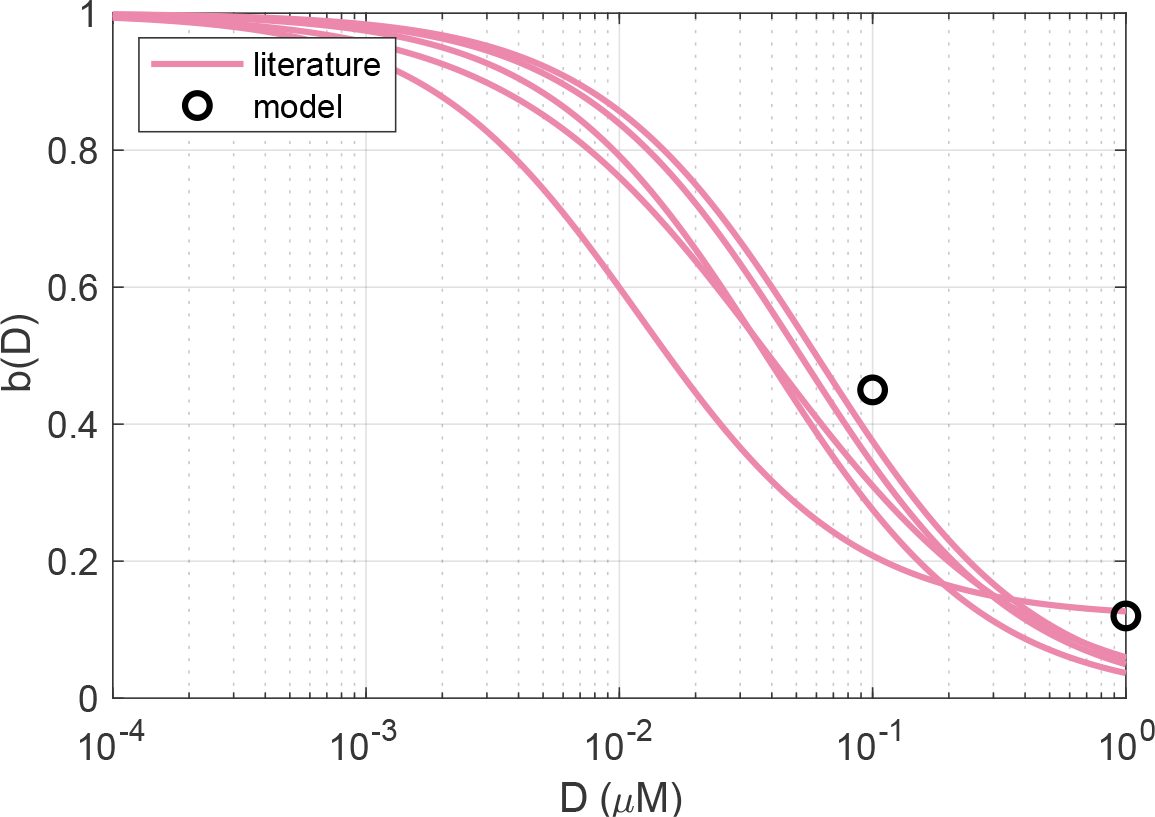
Comparison of the blocking effect of nifedipine *b*(*D*) used in our model to data from literature.

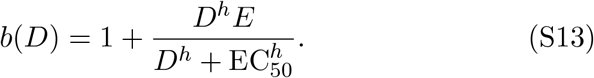

Here, as an example, *E* = *−*1 if the maximal effect of the drug is to block the current completely, and *E* = *−*0.5 if the maximal effect is to block the current by 50%. The values of EC_50_, *h*, and *E* used to generate the curves in Figure S2 are reported in Table S1. In cases where *h* or *E* are not provided, we have assumed *h* = 1 and *E* = *−*1.

**Table S1:** Dose-dependent effects of nifedipine on the L-type calcium current from literature used in Figure S2. The blocking effect is modeled using (S13).

| Reference | $\text{EC}_{50}$ | $h$ | $E$ |
| --- | --- | --- | --- |
| Kramer et al. 2013 [22] | $0.012 \mu\text{M}$ | 1.02 | -0.883 |
| Romero et al. 2018 [23] | $0.052 \mu\text{M}$ | | |
| DiStilo et al. 1988 [24] | $0.060 \mu\text{M}$ | | |
| Gibson et al. 2014 [25] | $0.039 \mu\text{M}$ | 0.85 | |
| Ma et al. 2011 [26] | $0.038 \mu\text{M}$ | | |

### S4 Model formulation

In this section, the formulation of the base model for the action potential of hiPSC-CMs, based on [10, 27] is provided. In the model formulation, the membrane potential (*v*) is given in units of mV, the Ca^2+^ concentrations are given in units of mM, all currents are expressed in units of A/F, and the Ca^2+^ fluxes are expressed as mmol/ms per total cell volume (i.e., in units of mM/ms). The parameters of the model are given in Tables S2–S8.

#### S4.1 Membrane potential

The membrane potential is governed by the equation

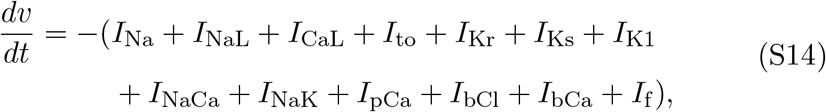

where *I*_stim_ is an applied stimulus current, and *I*_Na_, *I*_NaL_, *I*_CaL_, *I*_to_, *I*_Kr_, *I*_Ks_, *I*_K1_, *I*_NaCa_, *I*_NaK_, *I*_pCa_, *I*_bCl_, *I*_bCa_, and *I*_f_ are membrane currents specified below.

##### S4.1.1 Membrane currents

The currents through the voltage-gated ion channels on the cell membrane are in general given on the form

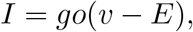

where *g* is the channel conductance, *v* is the membrane potential and *E* is the equilibrium potential of the channel. Furthermore, *o* = Π_*i*_ *z*_*i*_ is the open probability of the channels, where *z*_*i*_ are gating variables, either given as a function of the membrane potential or governed by equations of the form

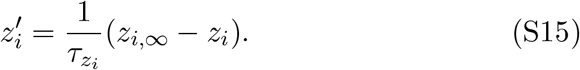

The parameters 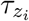 and *z*_*i,∞*_ are specified for each of the gating variables of the model in Table S9.

###### Fast sodium current

The formulation of the fast sodium current is an adjusted version of the model given in [28], supporting slower upstroke velocities more similar to those observed in the optical measurements of hiPSC-CMs. The current is given by

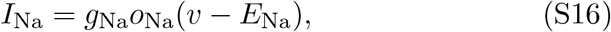

where the open probability is given by

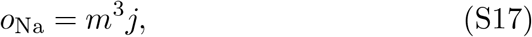

and *m* and *j* are gating variables governed by equations of the form (S15).

###### Late sodium current

The formulation of the late sodium current, *I*_NaL_, is based on [29] and is given by

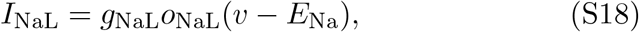

where the open probability is given by

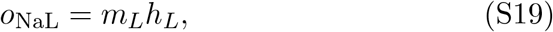

and *m*_*L*_ and *h*_*L*_ are gating variables governed by equations of the form (S15).

###### Transient outward potassium current

The formulation of the transient outward potassium current, *I*_to_, is based on [30] and is given by

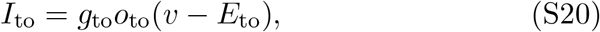

where the open probability is given by

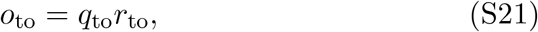

and *q*_to_ and *r*_to_ are gating variables governed by equations of the form (S15).

###### Rapidly activating potassium current

The formulation of the rapidly activating potassium current, *I*_Kr_, is based on [30] and is given by

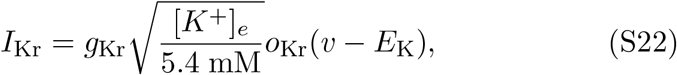

where

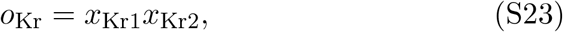

and the dynamics of *x*_Kr1_ and *x*_Kr2_ are governed by equations of the form (S15).

###### Slowly activating potassium current

The formulation of the slowly activating potassium current, *I*_Ks_, is based on [28] and is given by

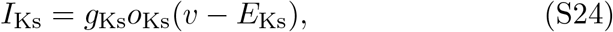

where

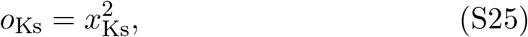

and the dynamics of *x*_Ks_ is governed by an equation of the form (S15).

###### Inward rectifier potassium

he formulation of the inward rectifier potassium current, *I*_K1_, is based on [28] and is given by

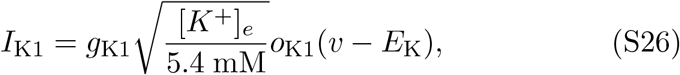

where

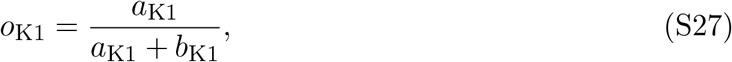

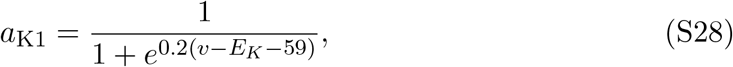

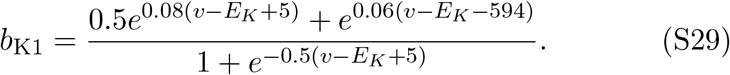

###### Hyperpolarization activated funny current

The formulation for the hyperpolarization activated funny current, *I*_f_, is based on [30] and is given by

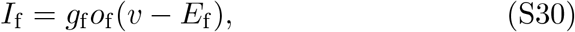

where

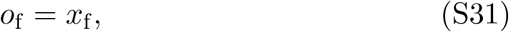

and the dynamics of *x*_f_ is governed by an equation of the form (S15).

###### L-type Ca^2+^ current

The formualtion for the L-type Ca^2+^ current, *I*_CaL_, is based on the formulation in [28] and is given by

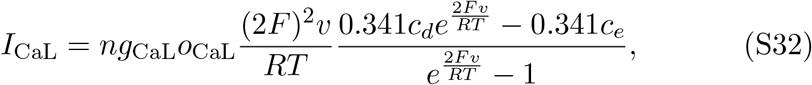

where

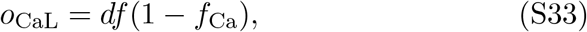

and the dynamics of *d, f* and *f*_Ca_ are governed by equations of the form (S15). Furthermore, *n* is the scaling factor for the number of L-type calcium channels in the cell membrane. In the paper, we consider two alternative models for this factor, one model directly based on [4, 8] given by (S9)–(S10) and one updated model described in the main paper and repeated in Section S4.4.

###### Background currents

The formulation of the background currents, *I*_bCa_ and *I*_bCl_, are based on [28] and are given by

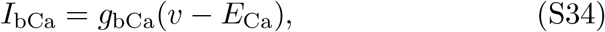

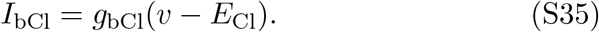

###### Sodium-calcium exchanger

The formulation of the Na^+^-Ca^2+^ exchanger current, *I*_NaCa_, is based on [28] and is given by

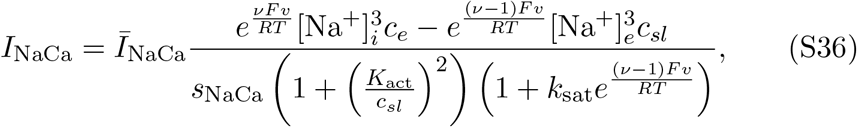

where

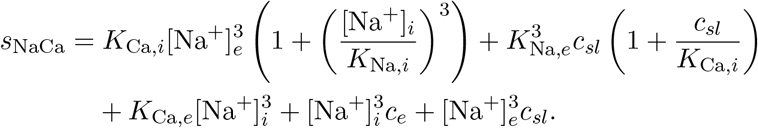

###### Sarcolemmal Ca^2+^ pump

The formulation of the current through the sarcolemmal Ca^2+^ pump, *I*_pCa_, is based on [28] and is given by

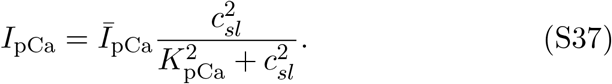

###### Sodium-potassium pump

The current through the Na^+^-K^+^ pump, *I*_NaK_, is based on [28] and is given by

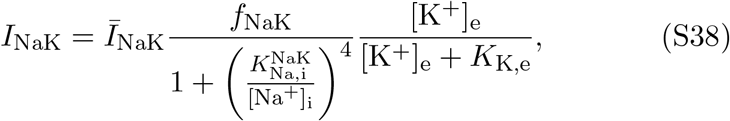

where

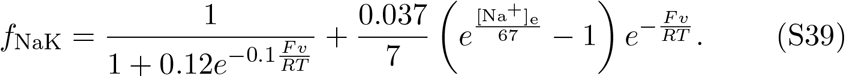

##### S4.1.2 Nernst equilibrium potentials

The Nernst equilibrium potentials for the ion channels are defined as

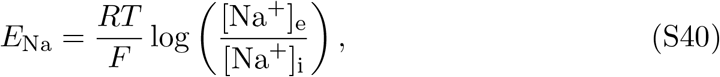

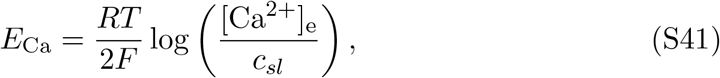

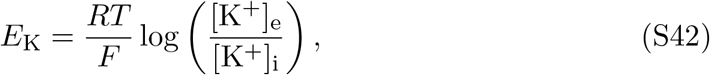

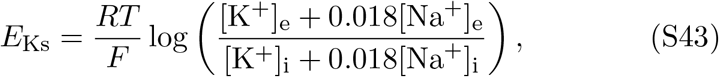

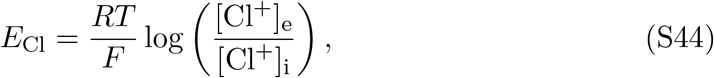

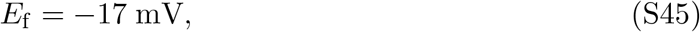

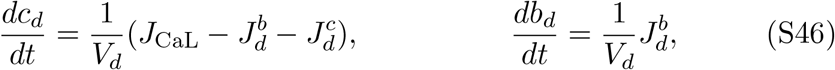

for the parameter values given in Table S3.

#### S4.2 Ca^2+^ dynamics

The Ca^2+^ dynamics are governed by

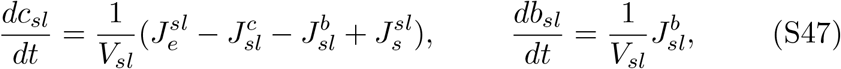

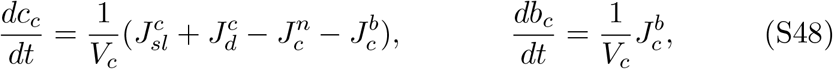

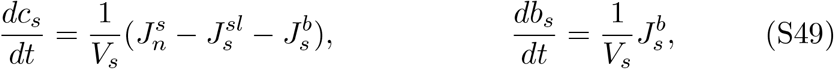

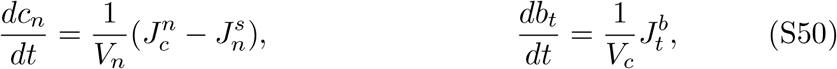

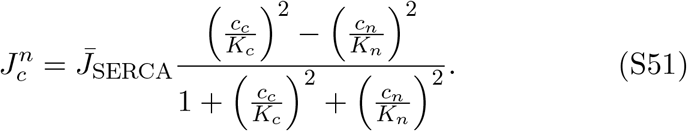

where *c*_*d*_ is the concentration of free Ca^2+^ in the dyad, *b*_*d*_ is the concentration of Ca^2+^ bound to a buffer in the dyad, *c*_*sl*_ is the concentration of free Ca^2+^ in the SL compartment, *b*_*sl*_ is the concentration of Ca^2+^ bound to a buffer in the SL compartment, *c*_*c*_ is the concentration of free Ca^2+^ in the bulk cytosol, *b*_*c*_ is the concentration of Ca^2+^ bound to a buffer that is not troponin in the bulk cytosol, *b*_*t*_ is the concentration of Ca^2+^ bound to troponin in the bulk cytosol, *c*_*s*_ is the concentration of free Ca^2+^ in the jSR, *b*_*s*_ is the concentration of Ca^2+^ bound to a buffer in the jSR, and *c*_*n*_ is the concentration of free Ca^2+^ in the nSR. The expressions for the fluxes are specified below.

##### S4.2.1 Ca^2+^ fluxes

###### Flux through the SERCA pumps

The flux from the bulk cytosol to the nSR through the SERCA pumps is given by

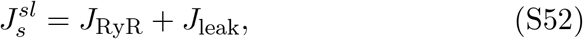

###### Flux through the RyRs

The flux from the jSR to the SL compartment is given by

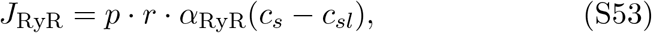

where *J*_RyR_ represents the flux through the active RyR channels and *J*_leak_ represents the flux through the RyR channels that are always open, given by

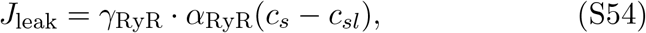

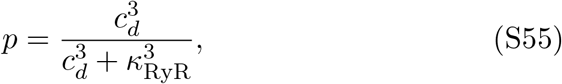

respectively. Here, *p* is the open probability of the active RyR channels given by

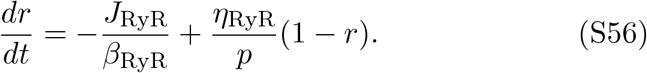

and *r* represents the fraction of RyR channels that are not inactivated and is governed by the equation

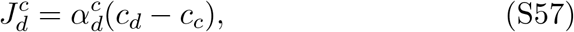

###### Passive diffusion fluxes between compartments

The passive diffusion fluxes between compartments are given by

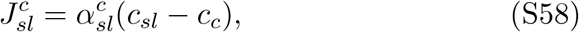

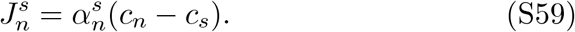

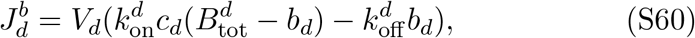

###### Buffer fluxes

The fluxes of free Ca^2+^ binding to a Ca^2+^ buffer are given by

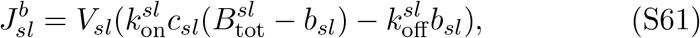

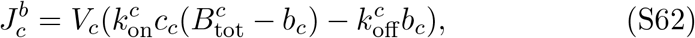

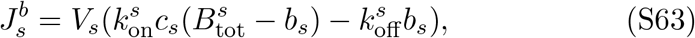

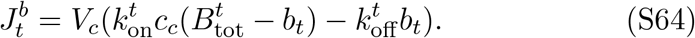

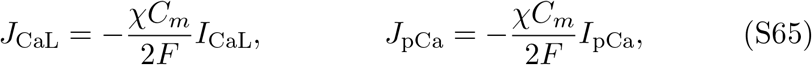

###### Membrane fluxes

The membrane fluxes, *J*_CaL_, *J*_bCa_, *J*_pCa_, and *J*_NaCa_, are given by

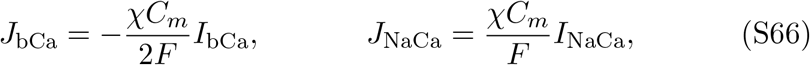

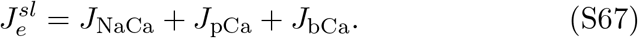

where *I*_CaL_, *I*_bCa_, *I*_pCa_, and *I*_NaCa_ are defined by the expressions given above. Furthermore,

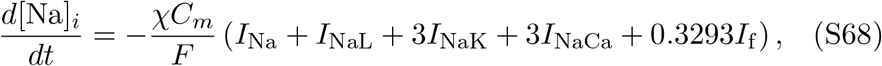

#### S4.3 Na^+^ dynamics

For the intracellular Na^+^ concentration, we use the same approach as in [27]. In this approach, spatial gradients of [Na^+^]_*i*_ in the cell are ignored and the concentration is governed by

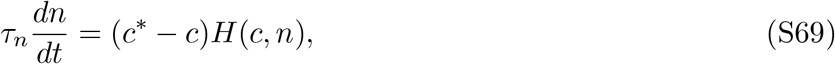

where the currents *I*_Na_, *I*_NaL_, *I*_NaK_, *I*_NaCa_, and *I*_f_ are specified above.

#### S4.4 Protein regulation

The scaling factor *n* for the number of L-type calcium channels on the cell membrane is modeled by

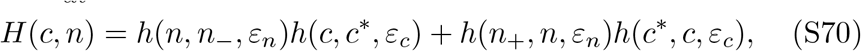

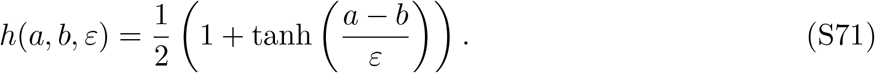

**Table S2:** Default geometry parameters of the base model.

| Parameter | Description | Value |
| --- | --- | --- |
| $V_d$ | Volume fraction of the dyadic subspace | 0.001 |
| $V_{sl}$ | Volume fraction of the SL compartment | 0.028 |
| $V_c$ | Volume fraction of the bulk cytosol | 0.917 |
| $V_s$ | Volume fraction of the jSR | 0.004 |
| $V_n$ | Volume fraction of the nSR | 0.05 |
| $\chi$ | Cell surface to volume ratio | $0.9 \mu\text{m}^{-1}$ |

**Table S3:** Physical constants and ionic concentrations of the base model.

| Parameter | Description | Value |
| --- | --- | --- |
| $C_m$ | Specific membrane capacitance | 0.01 pF/$\mu\text{m}^2$ |
| $F$ | Faraday's constant | 96.485 C/mmol |
| $R$ | Universal gas constant | 8.314 J/(mol·K) |
| $T$ | Temperature | 310 K |
| $[\text{Ca}^{2+}]_e$ | Extracellular $\text{Ca}^{2+}$ concentration | 1.8 mM |
| $[\text{Na}^+]_e$ | Extracellular sodium concentration | 155.3 mM |
| $[\text{K}^+]_e$ | Extracellular potassium concentration | 5.3 mM |
| $[\text{K}^+]_i$ | Intracellular potassium concentration | 59.5 mM |
| $[\text{Cl}^-]_e$ | Extracellular chloride concentration | 119.3 mM |
| $[\text{Cl}^-]_i$ | Intracellular chloride concentration | 15 mM |

**Table S4:**
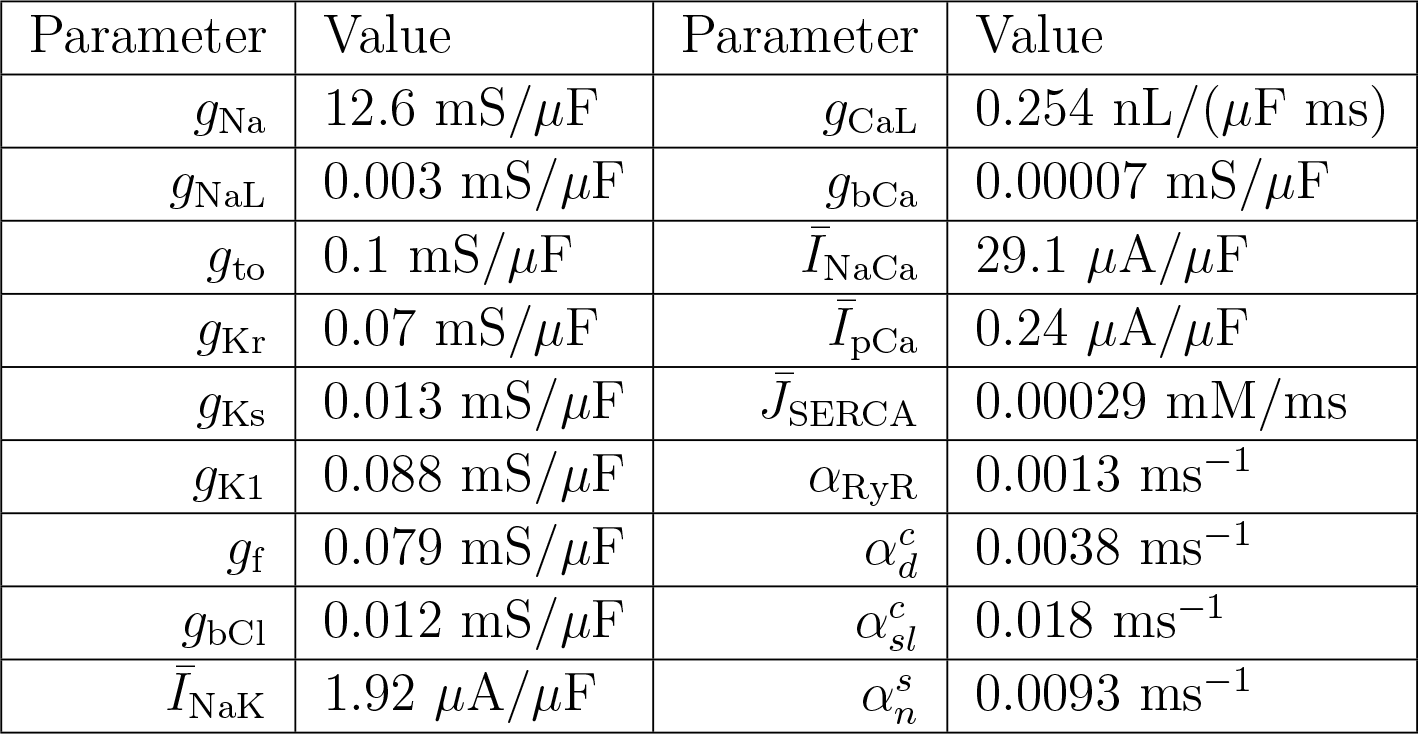
Conductance values and similar parameters for each of the membrane currents and intracellular Ca^2+^ fluxes of the base model.

**Table S5:** Parameters for the intracellular Ca^2+^ fluxes of the base model.

| Parameter | Flux | Value |
| --- | --- | --- |
| $K_c$ | $J_c^n$ | 0.00025 mM |
| $K_n$ | $J_c^n$ | 1.7 mM |
| $\beta_{\text{RyR}}$ | $J_s^{sl}$ | 0.027 mM |
| $\gamma_{\text{RyR}}$ | $J_s^{sl}$ | 0.001 |
| $\kappa_{\text{RyR}}$ | $J_{\text{RyR}}$ | 0.015 mM |
| $\eta_{\text{RyR}}$ | $J_s^{sl}$ | 0.00001 ms<sup>-1</sup> |

**Table S6:** Parameters for the membrane currents of the base model.

| Parameter | Current | Value |
| --- | --- | --- |
| $k_{\text{sat}}$ | $I_{\text{NaCa}}$ | 0.3 |
| $\nu$ | $I_{\text{NaCa}}$ | 0.3 |
| $K_{\text{act}}$ | $I_{\text{NaCa}}$ | 0.00015 mM |
| $K_{\text{Ca},i}$ | $I_{\text{NaCa}}$ | 0.0036 mM |
| $K_{\text{Ca},e}$ | $I_{\text{NaCa}}$ | 1.3 mM |
| $K_{\text{Na},i}$ | $I_{\text{NaCa}}$ | 12.3 mM |
| $K_{\text{Na},e}$ | $I_{\text{NaCa}}$ | 87.5 mM |
| $K_{\text{Na},i}^{\text{NaK}}$ | $I_{\text{NaK}}$ | 11 mM |
| $K_{\text{K},e}$ | $I_{\text{NaK}}$ | 1.5 mM |
| $K_{\text{pCa}}$ | $I_{\text{pCa}}$ | 0.0005 mM |

**Table S7:** Parameters for the Ca^2+^ buffers of the base model.

| Parameter | Compartment | Value |
| --- | --- | --- |
| $B_{\text{tot}}^c$ | Bulk cytosol | 0.034 mM |
| $k_{\text{on}}^c$ | Bulk cytosol | 40 ms<sup>-1</sup>mM<sup>-1</sup> |
| $k_{\text{off}}^c$ | Bulk cytosol | 0.03 ms<sup>-1</sup> |
| $B_{\text{tot}}^d$ | Dyad | 4.07 mM |
| $k_{\text{on}}^d$ | Dyad | 100 ms<sup>-1</sup>mM<sup>-1</sup> |
| $k_{\text{off}}^d$ | Dyad | 1 ms<sup>-1</sup> |
| $B_{\text{tot}}^{sl}$ | Subsarcolemmal space | 0.66 mM |
| $k_{\text{on}}^{sl}$ | Subsarcolemmal space | 100 ms<sup>-1</sup>mM<sup>-1</sup> |
| $k_{\text{off}}^{sl}$ | Subsarcolemmal space | 0.15 ms<sup>-1</sup> |
| $B_{\text{tot}}^s$ | Junctional SR | 27 mM |
| $k_{\text{on}}^s$ | Junctional SR | 100 ms<sup>-1</sup>mM<sup>-1</sup> |
| $k_{\text{off}}^s$ | Junctional SR | 39.6 ms<sup>-1</sup> |

**Table S8:** Parameters for the regulation of the number of L-type calcium channels in the cell membrane.

| Parameter | Value | Parameter | Value |
| --- | --- | --- | --- |
| $\tau_n$ | 400 mMms | $c^*$ | 5.39·10<sup>-5</sup> mM |
| $n_-$ | 0.1 | $n_+$ | 3 |
| $\varepsilon_n$ | 0.01 | $\varepsilon_c$ | 10<sup>-7</sup> mM |

**Table S9:**
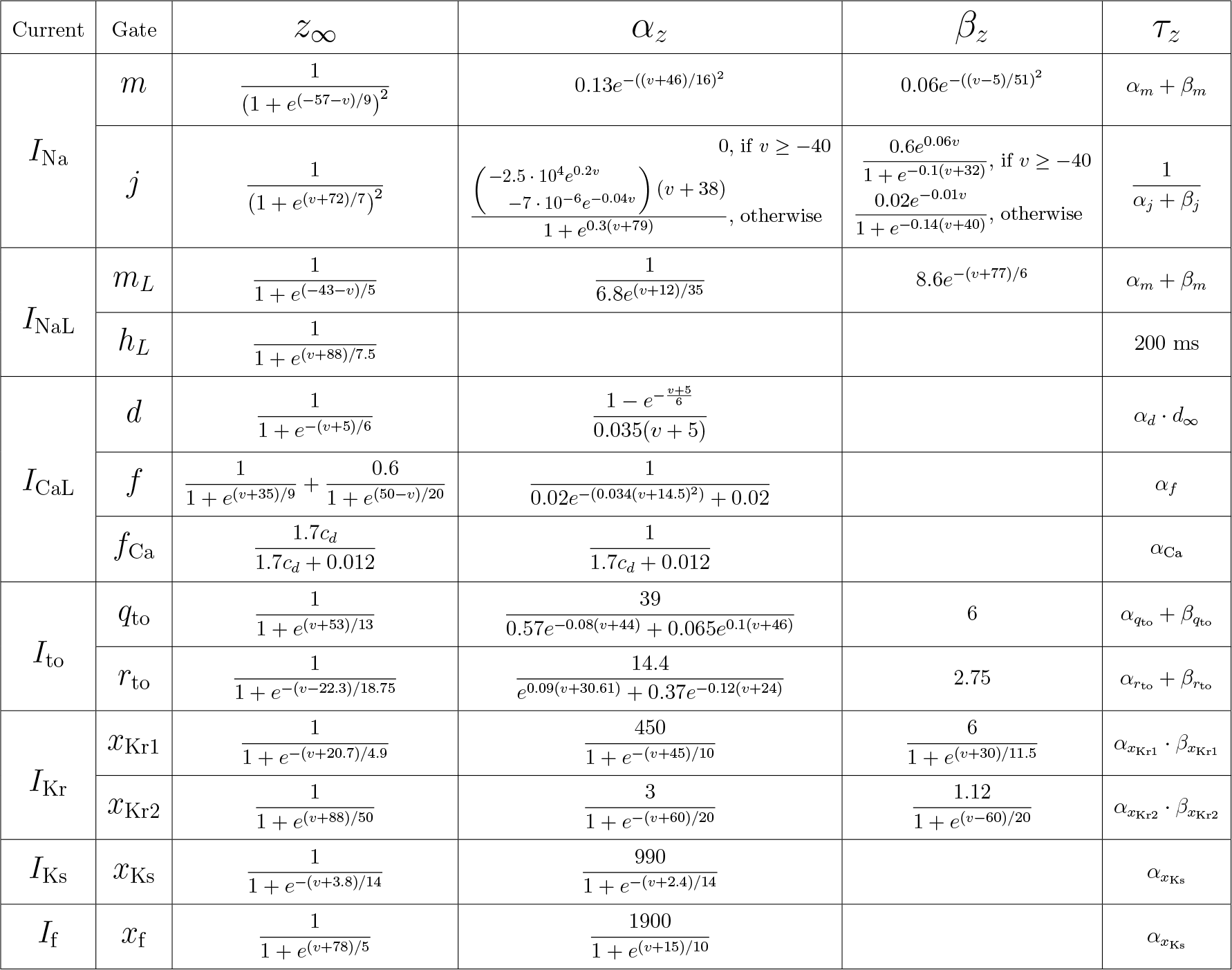
Specification of the parameters *z*_*∞*_ and *τ*_*z*_, for *z* = *m, j, m*_*L*_, *h*_*L*_, *d, f, f*_Ca_, *q*_to_, *r*_to_, *x*_Kr1_, *x*_Kr2_, *x*_Ks_ and *x*_f_ in the equations for the gating variables (S15).

## Footnotes

1 Note that we here use the notation *r* for the variable that is called *m* in [4, 8]. The reason is to avoid confusion with the variable *m* in the formulation of the model used in this study.

